# Binding items to contexts through conjunctive neural representations with the Method of Loci

**DOI:** 10.1101/2024.12.19.629352

**Authors:** Jiawen Huang, Akshay Manglik, Nick Dutra, Hannah Tarder-Stoll, Taylor Chamberlain, Robert Ajemian, Qiong Zhang, Kenneth A. Norman, Christopher Baldassano

## Abstract

Schematic prior knowledge can provide a powerful scaffold for episodic memories, yet the neural mechanisms underlying this scaffolding process are still poorly understood. A crucial step of the scaffolding process is the way in which details of a new episode are connected to an existing schema, forming a robust memory representation that can be easily accessed in the future. A unique testbed for studying this binding process is a mnemonic technique called the Method of Loci (MoL), in which people meaningfully connect items to be remembered with a well-learned list of imagined loci. We collected fMRI data from participants in 3 longitudinal sessions while they were enrolled in a month-long MoL training course, all of whom showed dramatic improvements in their ability to remember lists of 20 or 40 words. We obtained neural patterns when the loci and objects are presented by themselves, when they are combined into an integrated representation at encoding, and when the integrated representation was subsequently retrieved, as well as verbal descriptions from the participants about the way they associated each item to each locus. We found that in default mode network regions, including medial prefrontal cortex (mPFC), the combined representations of loci and items are highly conjunctive: the unified locus-item representation was substantially different from a linear combination of the isolated locus and item representation, reflecting the addition of new integrative details specific to each combined pair. The conjunctive component of the representation reflected the particular creative details generated by individual participants and increased over time as participants gained expertise in MoL. Our findings reveal a critical role for the default mode network in creating meaningful connections between new information and well-learned schematic contexts.

## Introduction

Decades of research on human memory have sought to probe the functional architecture of the memory system by using memorization tasks with word lists and pictures (Howard & Kahana, 2002; Polyn et al., 2009; Puff, 1979). Paradoxically, participants often struggle to recall these kinds of simple items (Murdock, 1974), while showing impressive ability to remember much more complex stimuli such as movie events (Chen et al., 2017). Stimuli drawn from familiar real-world settings allow us to draw on *schemas,* our prior knowledge about the structure of the world and how events unfold over time. This prior knowledge can scaffold memory processes in various ways. Past behavioral research has studied how schemas facilitate memory during the encoding process (e.g., Bartlett, 1932; Chase & Simon, 1973; Gasser & Davachi, 2023; Gobet & Waters, 2003; Huang, Furness, et al., 2023; Masís-Obando et al., 2024) and how schemas aid memory through providing a scaffold at retrieval (e.g., J. R. Anderson, 1981; R. C. Anderson & Pichert, 1978; Huang, Velarde, et al., 2023; Watkins & Gardiner, 1979). For this scaffolding to be effective, the details to be remembered need to be “attached” to the schema; that is, there must be a meaningful relationship between the current episode and schematic knowledge that is formed during encoding and accessible during retrieval. Despite extensive past research, this crucial process of how event-specific details are combined with schemas to form a robust memory representation remains relatively unexplored. The main aim of the current study is to investigate the neural mechanism behind this binding of schemas and event details.

Past research on associative memory has shown that effectively linking items to an externally-presented context requires more than simply experiencing the item in that context; the item must also interact with the context in a meaningful way (Eich, 1985; Murnane et al., 1999; Shin et al., 2021). This should also be true when the context arises internally through the activation of structured knowledge, but it is difficult to study this process in realistic events.

Schemas and event details are often tightly intertwined (e.g., a metal detector is closely linked to an airport schema), and this inherent integration makes it challenging to disentangle the details from the schema and understand how they are combined in memory. Thus, even though recent neuroimaging studies have highlighted the role of the Default Mode Network (DMN) regions in representing schemas (Baldassano et al., 2018; De Soares et al., 2024; Gilboa & Marlatte, 2017; Masís-Obando et al., 2021; van Kesteren et al., 2012; Van Kesteren et al., 2013), the neural mechanisms underlying the interaction between schemas and details in an ongoing event are still unclear.

We hypothesized that when schemas and event details are interactively combined together, the resulting representation should be conjunctive (O’Reilly & Rudy, 2001), meaning that the memory formed is more than a linear sum of its constituent components. For example, when a chess grandmaster sees a chess board, they build a mental representation that is not simply a list of the pieces and their positions, but also includes the relationships between the pieces and the potential dangers and opportunities afforded by these relationships. Some previous work with fMRI has shown signatures of conjunctive representations when perceiving individual objects created from simple features (Erez et al., 2016; Liang et al., 2020), images of human-object interaction (Baldassano et al., 2017), or scenes with different features of environment, objects, and people (van den Honert et al., 2017). While these studies examined conjunctive representations during perception, here we investigated the role of conjunctive representations in linking contexts and items to form a robust episodic memory that can be reinstated through schema-based retrieval.

We developed a new paradigm for studying conjunctive representations based on an ancient mnemonic technique called the Method of Loci (MoL, Figure 1a). The technique involves building and consolidating a spatial layout (memory palace) in the mind, with ordered locations (loci) in the memory palace that serve as a schematic scaffold. During encoding, each item to be remembered is combined with each of the loci in order by forming a meaningful connection between the two, such as imagining an event involving that item occuring at that locus. During retrieval, people mentally retrace their steps through the memory palace in order, using each locus as a cue to recall its associated item. A critical skill for using this technique is the ability to create and elaborate on a relationship between the item and locus, since adding a meaningful interaction is highly effective at improving associative memory (reviewed in Higbee, 1979). Mnemonists will in fact strategically choose loci with high “associability” (i.e. that have a wide range of features, attributes, and associations) in order to facilitate the creation of these interactions (Bellezza, 1996).

**Figure 1.**
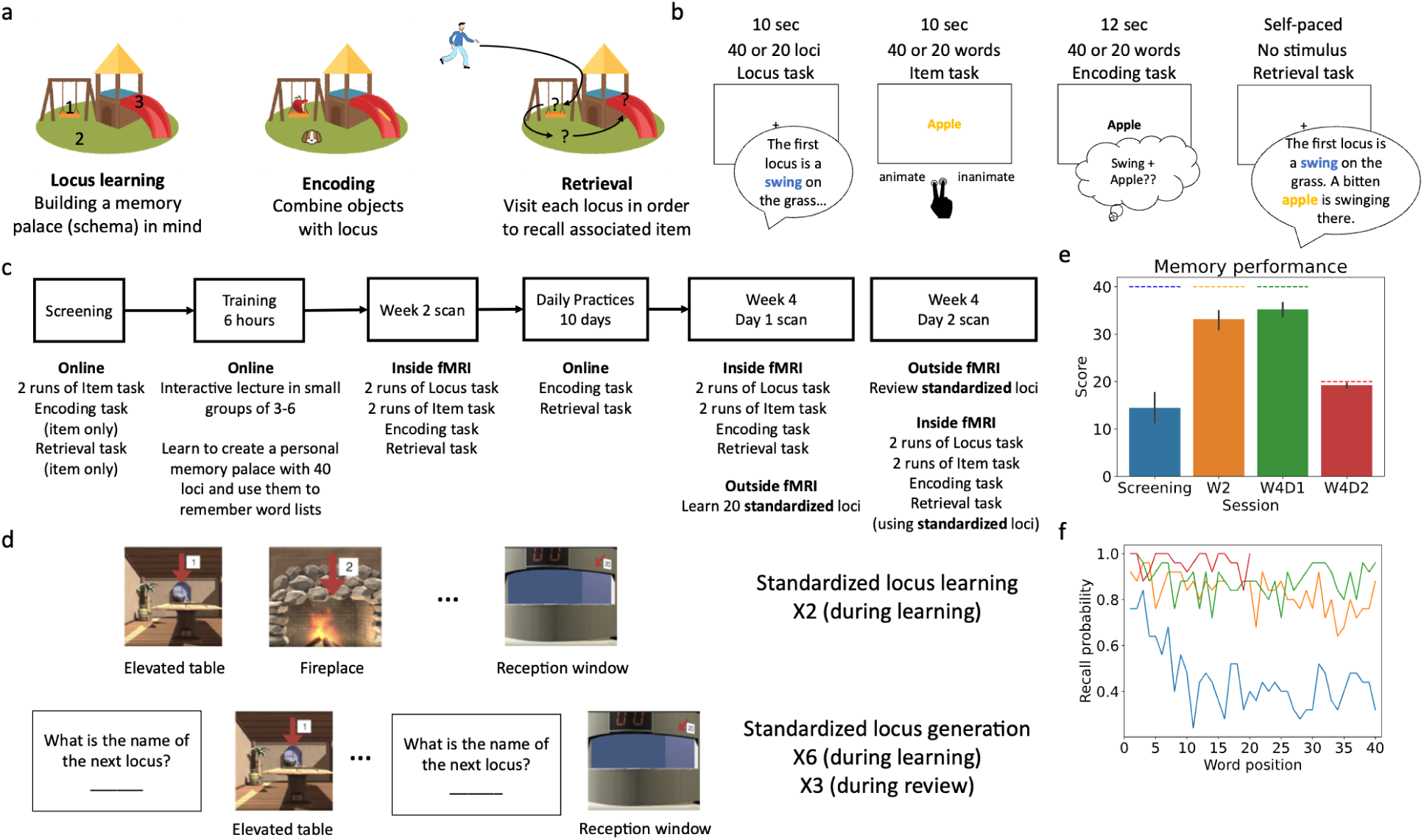
Illustration of the paradigm and behavioral performance. **a.** Demonstration of the Method of Loci, in which items to be remembered are attached to sequential locations along an imagined spatial map. **b.** The tasks participants completed in the scanner, capturing: neural representations of loci alone, items alone, encoding an item at each locus, and retrieving an item at each locus. **c.** Illustration of the four-week training and data collection schedule for participants. **d.** Illustration of how participants learned the standardized loci. **e.** Memory performance (number of words correctly recalled in order) for participants at different timepoints during the study. Dashed lines represent the maximum score possible in each session. Error bars represent the 95% confidence interval. Participants demonstrated substantial improvement after receiving two weeks of training, continued to improve at week 4, and were able to generalize to the standardized loci on W4D2 with near-ceiling accuracy. **f.** Probability of recalling a word given the word’s position in the encoding list. During screening (blue), participants showed a strong primacy effect (they were much more likely to recall words in the beginning of the list). However, in later sessions (W2: orange, W4D1: green, W4D2: red) this effect was greatly reduced.

While some past research has studied MoL using brain imaging, this work has largely focused on the technique itself and how it changes overall patterns of brain activity (C. Liu et al., 2022; Maguire et al., 2003; Wagner et al., 2021). In one of the first neuroimaging works on MoL, Maguire et al. (2003) showed increased hippocampal activity while memory experts remembered information using MoL. More recent research has focused on fMRI functional connectivity differences between experts and novices, finding that connectivity in novices becomes more similar to experts’ after training (Wagner et al., 2021). In the current study, we use MoL as a testbed for studying how novel information is encoded into a well-consolidated schematic map, by measuring the neural representations of each item and locus on their own and then relating these to neural patterns found at encoding (when the item and locus are being bound together) and retrieval.

We recruited participants for a 4-week MoL training program that drastically improved their memory for word lists, and collected fMRI data in one session after week 2 and in two sessions after week 4. In the scanner, participants described their loci and imagined words on the screen. They also used the MoL technique to encode a list of words by combining each word with a locus in their memory palace. Finally, they recalled the list of words by describing each locus, item, and how they were combined together. We used multivariate analyses to investigate how loci and items are represented separately (in locus description and item imagination) and combined (in encoding and retrieval) during MoL. We hypothesized that the activity during encoding and retrieval should be conjunctive, i.e. that it contains information that goes beyond a linear combination of the locus and item. We measured the conjunctive representations in multivariate neural activity patterns (the component of neural activity not predictable from the locus pattern and item pattern) and also assessed the amount of conjunctivity in semantic space (the degree to which the verbal descriptions given by the participants incorporated semantic content beyond that of the locus alone and item alone). We hypothesized that the amount of conjunctive representation in both neural and semantic spaces should increase over the course of the training as participants developed expertise in the technique. Additionally, the two measures of conjunctive representation should be related to each other, with the degree of neural conjunctivity for a locus-item pair predicting the degree of semantic conjunctivity for that pair.

Past research on conjunctive representations has focused on the role of the hippocampus in forming conjunctive representations to associate two novel, arbitrary, and independent events in memory (Squire et al., 1989). However, forming conjunctive representations of real-life events supported by schemas might require involvement of the cortex. Previous research has proposed that remembering details consistent with a schema relies on cortical regions like medial prefrontal cortex (Brod et al., 2015; Ghosh & Gilboa, 2014; Gilboa & Marlatte, 2017; Z.-X. Liu et al., 2017; Preston & Eichenbaum, 2013; Raykov et al., 2020, 2021; Reagh & Ranganath, 2023; Tse et al., 2007; van Kesteren et al., 2012; Van Kesteren et al., 2013; van Kesteren et al., 2020), potentially because episodic memory can be consolidated rapidly through interaction with the activated associative schema (Morris, 2006). The MoL provides a way to make *any* item schema-consistent, by having participants imagine a situation or find a dimension in which the item is connected to the locus (generating additional details as necessary to reinforce the schematic relationship they identified). In addition, studies have found representations of naturalistic schemas (Baldassano et al., 2018; De Soares et al., 2024; Reagh & Ranganath, 2023) in the DMN. Thus, we hypothesized that we would find conjunctive representations of locus-item pairs in DMN regions, including medial prefrontal cortex (mPFC), which has been shown to be most sensitive to the top-down internal activation of naturalistic schemas (De Soares et al., 2024). In line with this prediction, we found that conjunctive representations were present through the DMN (and beyond) – memory representations were in fact *dominated* by this conjunctive information rather than pure locus or item representations. The amount of conjunctive representation increased in the novices over the course of training, and was related to the amount of additional creative details added to the story that go beyond descriptions of the locus and item on their own. Overall, these findings point to a crucial role of the conjunctive representation in the DMN for MoL, and the importance of DMN more broadly in forming conjunctive associations that support robust memory.

## Results

An overview of the four-week study paradigm is presented in Figure 1c. After passing an online screening, novice participants (N = 25) first participated in an online training program for 2 weeks, in which they were taught the MoL technique and created a memory palace consisting of 40 loci. During the training program, participants were also familiarized with the 40 items (20 animate, 20 inanimate) that they would be associating with the loci in subsequent phases of the study. Participants were given their first fMRI scan in week 2 (W2), which consisted of four types of tasks (Figure 1b). In the locus task (repeated twice), participants verbally described each of their loci for 10 sec, before being prompted to describe the next locus. In the item task (repeated twice), participants saw a word for 10 sec and were asked to imagine the word vividly and then make a judgment of whether or not the word was a living object. In the encoding task, participants used MoL to encode each word for 12 sec, by forming an association between the item and the locus in the memory palace. In the retrieval task, participants described each locus in their memory palace, the item that the locus was connected to, and the story they created to link the locus to the item. The recall was self-paced, but they were encouraged to spend at least 10 sec for each pair. After the first scan, participants completed 10 daily online practices for two more weeks, in which they were presented with 40 words to remember using MoL, each for 12 sec, and then attempted to recall the words in the correct order. In week 4, they were scanned on two consecutive days. On week 4 day 1 (W4D1), they completed a scan just like the week 2 scan. After that, they were taught (outside of the scanner) a new, standardized memory palace with 20 loci (Figure 1d), presented as videos on a laptop. On week 4 day 2 (W4D2), they reviewed the standardized loci and then completed a scan with the same structure as the previous two, now using the standardized loci to remember 20 words (a subset of the 40 words used previously). Note that the same 20 words were presented in the same order to all the participants, so they were each attempting to create a binding between the same set of item-locus pairs. Participants’ performance in the memory task showed a massive improvement compared to the baseline (Figure 1e), from 14.44 (SD = 7.93) at baseline to 33.16 (SD = 5.07) to 35.24 (SD = 3.72) in Week 2 and Week 4 respectively. Participants were also near ceiling on the standardized memory task, with a mean score of 19.2 (SD = 1.02) out of 20 words in total. Participants’ serial position curves were markedly different after training (Figure 1f), shifting from showing a standard primacy effect (Murdock, 1974) to near-uniform recall of words from all serial positions.

### Widespread representations of locus and item by themselves and during encoding

Using a generalized linear model (GLM), we obtained the representation of each locus and item alone in each run of the locus and item task and the representation of the locus-item pair during encoding and retrieval (Figure 2a). We then measured the pattern correlation (within searchlights on the cortical surface) between corresponding loci or items in the two runs of the locus and item tasks (while accounting for overall task similarity not specific to particular loci or items; see Methods). For similarity between locus/item and encoding, we used a similar approach, measuring the pattern correlation between the locus by itself (e.g., “swing”) or item by itself (e.g., “apple”) with the locus-item (e.g., “swing-apple”) pair where the locus/item was used.

**Figure 2.**
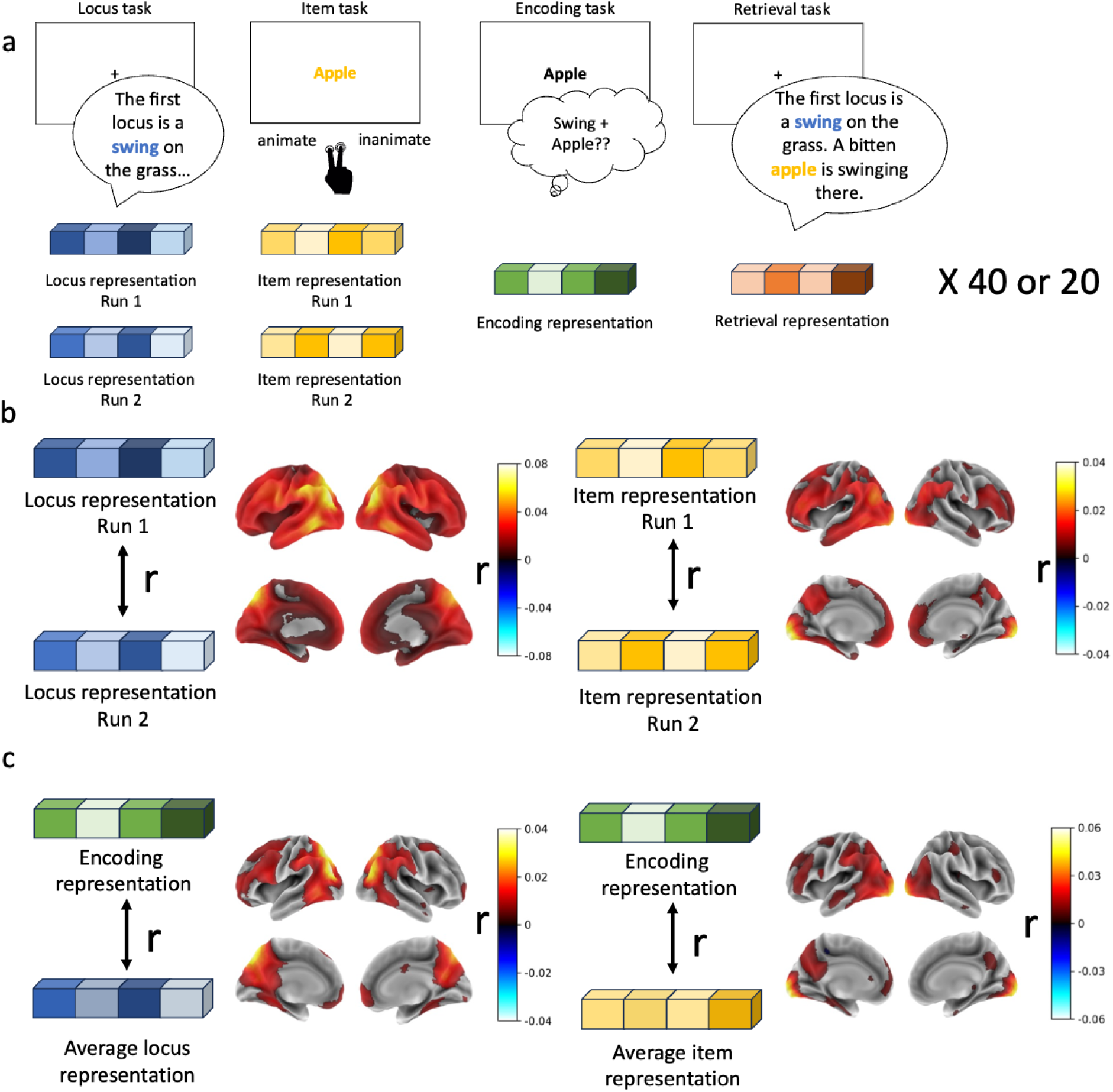
Brain regions representing loci and items alone and during encoding. **a.** Illustration of how representations were obtained. For each searchlight on the cortical surface, the multivariate activity pattern was measured for each locus (in each two runs), item (in each of two runs), and locus-item pair during encoding and retrieval of 40 words (in the first two sessions) or 20 words (in the final session). **b.** Representation of locus and item in the brain. Almost the whole brain showed significant locus-specific activation (pattern similarity for corresponding loci in the two locus runs), with the strongest effects in angular gyrus (AG) and posterior medial cortex (PMC). For two item runs, visual cortex, AG, PMC, and mPFC all showed item-specific activation patterns. **c.** Representation of locus and item during encoding. The current (imagined) locus was represented in AG, PMC, and mPFC during encoding, while items (presented as words) were represented in visual cortex and PMC. The brain maps were thresholded based on results of a permutation test, at q < .05 with FDR correction.

We showed that a large portion of the brain represented imagined loci, most notably in angular gyrus (AG) and posterior medial cortex (PMC) (Figure 2b, left). The same analysis for the item task (during which each item was presented as written word) showeditem representation in visual cortex, AG, PMC, and mPFC (Figure 2b, right). Because using MoL requires a combination of locus and item during encoding, we looked for representations of locus and item information during encoding (Figure 2c). Locus reactivation was observed in the default mode network regions, including AG, PMC, and mPFC, while item information was represented primarily in the visual cortex and PMC. These results demonstrate that participants were successfully reinstating locus-specific patterns throughout a broad network of regions during encoding, while simultaneously maintaining a representation of the presented item to be remembered.

### Encoding residuals track idiosyncratic semantic combinations of loci and items

When people use MoL to remember a word, in addition to thinking simultaneously about the item and the locus, they also add in additional details to forge a link between the two, creating a conjunctive representation between the locus and the item. We therefore sought to measure the conjunctive component of the encoding and retrieval representations by decomposing them into a representation of the locus, the item, and the conjunction between the locus and the item. For each locus-item pair (separately for each participant and brain region), we regressed out the representation of locus and item from encoding (Figure 3a), yielding an *encoding residual* containing the conjunctive information (the pattern for this locus-item pair that is not linearly related to the isolated locus and isolated item patterns) but also potentially including representations of irrelevant (e.g. off-task) mental state and measurement noise not reflective of neural activity. One way to test if the encoding residual contains meaningful conjunctive information (vs noise) is to check whether it is related to the semantic content of participants’ verbal recall. We focused on the W4D2 session, in which participants used the same memory palace to remember the same set of words in the same order, but often came up with very different ways of relating a locus to an item (see Figure 3b, rightmost column for examples). In three key regions of interest in the DMN, found in previous studies to represent schematic knowledge (Figure 3c, Baldassano et al., 2018; Yeo et al., 2011), we investigated whether higher similarity of the encoding residual representations in these regions correlates with semantic similarity in participants’ verbal descriptions. For all the ROI analyses presented below, we also conducted the same analyses in anterior, posterior, and the whole hippocampus and found no significant effects in all of the analyses (Figure S1). For each locus-item pair, we constructed a between-subject representational similarity matrix of the (neural) encoding residual (Kriegeskorte et al., 2008); we also computed a semantic similarity matrix based on people’s verbal recall of the locus-item pair using a cross-encoder with sentence-BERT (Reimers & Gurevych, 2019) (Figure 3b); we then computed the Spearman correlation between the lower triangles of these matrices, and compared this to a null distribution. In AG and mPFC, if two people showed similar neural patterns in the encoding residual, they were also more likely to tell similar stories (AG: z = 3.75, p = .001; mPFC: z = 2.32, p = .017), though this effect was not found for PMC (z = −0.96, p = .349) (Figure 4d). This demonstrates that the encoding residual representations in the DMN indeed track idiosyncratic content used to link a locus to an item, supporting the idea that they contain information about the conjunction of the locus and the item.

**Figure 3.**
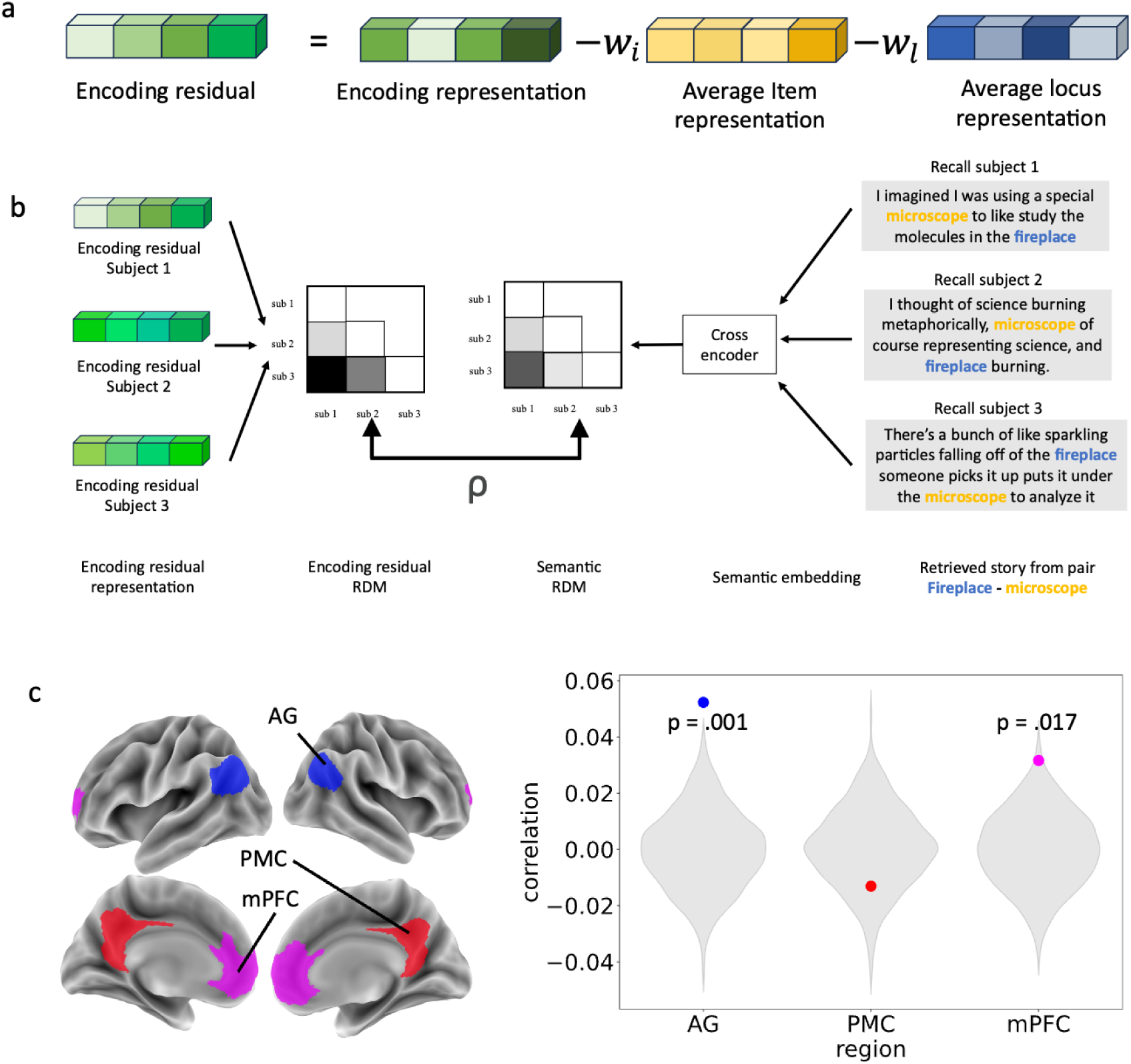
Encoding residuals track semantic similarity across stories. **a.** Illustration of how the encoding residual is computed. Locus and item representations were removed from the encoding representation via linear regression, and the resulting encoding residual contained the conjunctive representation of the linkage between locus and item. **b.** Illustration of the RSA analyses. The analysis is based on week 4 day 2, when participants used the same memory palace to remember the same words, allowing us to see the idiosyncratic item-in-locus story each person generated. Pairs of recalls from different subjects for the same story were input to a cross-encoder language model, generating a semantic representational similarity matrix quantifying the semantic similarity between each pair of stories. Similarly, the encoding residuals were compared across pairs of participants to create a neural representational similarity matrix. We then looked at the similarity of the neural representational similarity matrix to the semantic representational similarity matrix. **c**. Demonstration of location of the ROIs on the cortical surface. **d.** Neural-semantic correlation in the three ROIs. The grey violin plot represents the null distribution generated from shuffling the subject correspondence between the two measures. The dots show the true correlation between the neural representational similarity matrices and semantic representational similarity matrices. In AG and mPFC, similar neural representations track similar semantic representations of stories.

**Figure 4.**
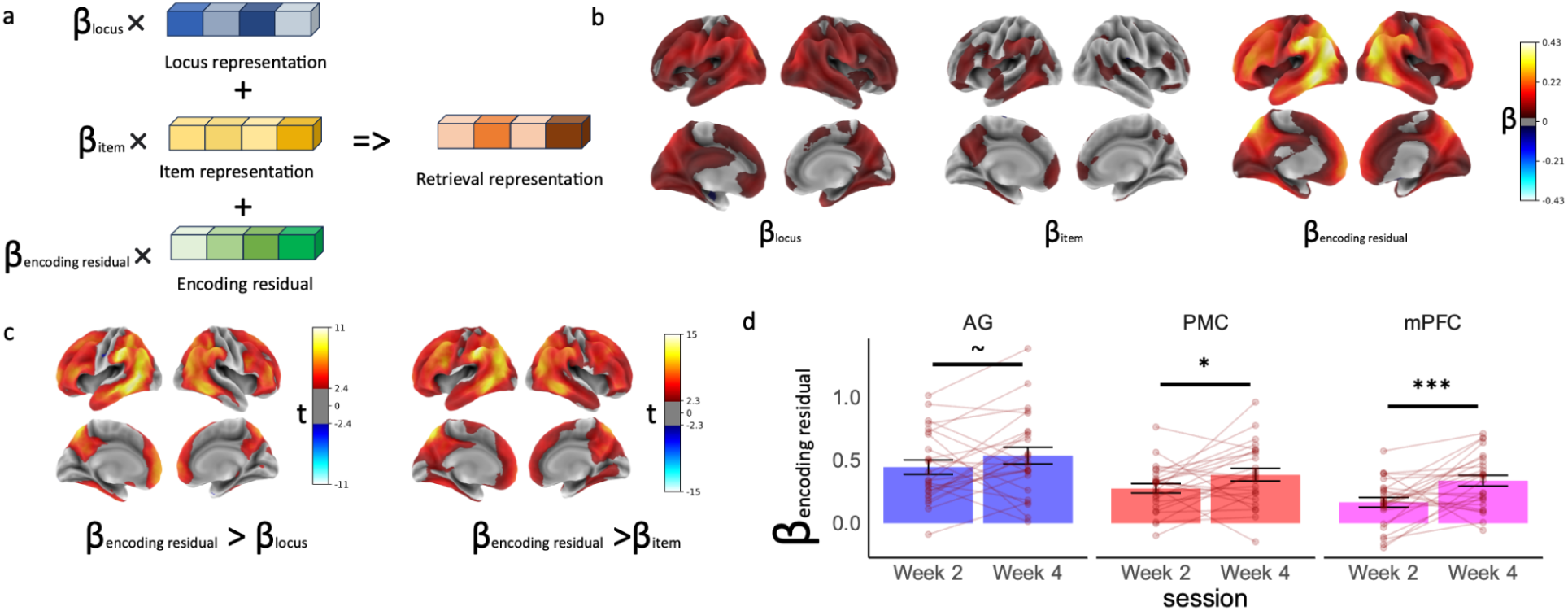
Conjunctive representation in the brain. **a.** Illustration of how the strength of the conjunctive representation was measured. When using locus, item, and encoding residual to predict retrieval representation, the weight of the encoding residual indicated the amount of conjunctive representation. **b.** The weight for locus, item, and encoding residual when predicting recall patterns in each searchlight. Widespread and robust representations were found for locus and the encoding residual during retrieval, with the strongest locus effects in AG and PMC and the strongest residual (conjunctive) effects in AG, PMC, and mPFC. Item representations were also found in AG, PMC, and mPFC, though with a more limited overall extent of significant effects. **c.** Comparisons between weights for encoding residual and locus (left) and encoding residual and item (right). Encoding residual was more strongly reinstated than either locus or item, especially in AG, PMC, and mPFC. In b and c, color indicates significant difference at q<.05 with FDR correction. **d.** Weight of encoding residual in week 2 and week 4 in the three ROIs. Error bars represent standard error of the mean. Points and connections represent individual participants. All ROIs had significantly positive weights for the encoding residual and the weight of encoding residual increased from week 2 to week 4 in all ROIs (∼ p =0.08, * p < .05, *** p < .001).

### Widespread and robust conjunctive representation in the brain

The encoding residual contains not just the conjunctive representation, but also irrelevant representations and noise, making it difficult to estimate the strength of the conjunctive representation based solely on the norm of the encoding residual. We can instead look at the extent to which the encoding residual is reinstated at recall: only the conjunctive component of the residual should be reactivated when retrieving the memory using the locus as a cue. For each locus-item pair (separately for each participant and brain region), we ran a regression to predict retrieval representation from the locus representation, item representation, and encoding residual representation (Figure 4a, bottom). To ensure that these regression weights were specific to the locus-item pair (rather than reflecting a generic task-related representation), the weights were adjusted relative to a null distribution created by permuting which retrieval pattern was matched to the item, locus, and encoding residual patterns (see Methods). Importantly, the weights for the encoding residual capture the amount of conjunctive representation for the pair.

For the locus and item weights in the retrieval regression, we found similar results to the locus-encoding correlation and item-encoding correlation described above, with locus representations in AG, PMC, and item representations in AG, PMC, and mPFC during retrieval (Figure 3b). Strikingly, we found that the encoding residual is represented very strongly in AG, PMC, and mPFC during retrieval, showing the importance of the conjunctive representation during the MoL. We conducted a paired t-test to compare the weight of the encoding residual to the weight of the locus and item, and showed that in largely overlapping regions including AG, PMC and mPFC, the encoding residual was represented more strongly than the locus or item by themselves (Figure 3c).

### Relationship between conjunctive representation in the DMN and training and behavior

We next explored whether the conjunctive representation was related to expertise and recall behavior. Given the importance of the conjunctive representation to neural representations and the centrality of locus-item binding in the technique of MoL, we would expect differences in the amount of conjunctive representations over the course of training. Comparing each subject’s average weight for the encoding residual (across items) against 0, we found that three ROIs (AG, PMC, and mPFC) demonstrated a significant weight of encoding residual in both week 2 and week 4 (all p < .001), with a higher weight in AG than mPFC (t = 6.58, p < .001) and PMC (t = 5.39, p < .001) and a higher weight in PMC than mPFC (t = 2.71, p = .009). To look at the change in weight of the encoding residual with expertise, we used a mixed-effects linear model to predict the residual weight from stage of training (week 2 vs. week 4) while controlling for the duration of recall (to account for the possibility of increased encoding-recall similarity due solely to retrieval pattern estimates being more stable for longer recalls) with a random subject intercept. We found that, across all ROIs, there was an increase in the weight of encoding residual from week 2 to week 4 (Figure 4b), which was significant in PMC and mPFC (PMC: t = 2.24, p = .025; mPFC: t = 3.77, p < .001) and marginally significant in AG (t = 1.88, p = .06), providing evidence that representations become more conjunctive with increasing expertise in MoL. As a control, we ran similar linear mixed-effect regressions to test whether the contribution of locus or item representations to either encoding or retrieval representations varied with training, and did not find any significant effects of expertise (all p > .131).

We conceptualized conjunctive representation as the additional details participants added for linking the locus to the item, which in the neural space is measured as the weight of encoding residual, taking into account the locus and item representations. We can define a similar measure for verbal recall, reflecting the extent to which the individual stories generated by each person added new semantic content not present in the locus or item alone. We quantified this by computing the semantic distance of the story to the locus-item pair using the cross-encoder described above (where the semantic distance is equal to 1 minus the similarity from the cross-encoder) – we refer to this measure as *story deviation*. If this story deviation measure is high, the generated story is more different from just the locus and item pair (Figure 5a), as can be observed in example stories with high and low story deviations (Figure 5b). Comparing week 2 and week 4, we conducted a linear mixed-effects regression predicting story deviation from session, with a random subject intercept. We found a significant increase in story deviation across sessions (t = 2.71, p = .007) (Figure 5c). Across subjects, increase in story deviation between week 2 and week 4 is positively correlated with increase in the weight of encoding residual in all the ROIs (AG: r = .402, p = .046; PMC: r = .540, p = .005; mPFC: r = .463, p = .020). These results provide converging evidence to suggest that forming a good conjunctive representation is a skill that is associated with training of MoL.

**Figure 5.**
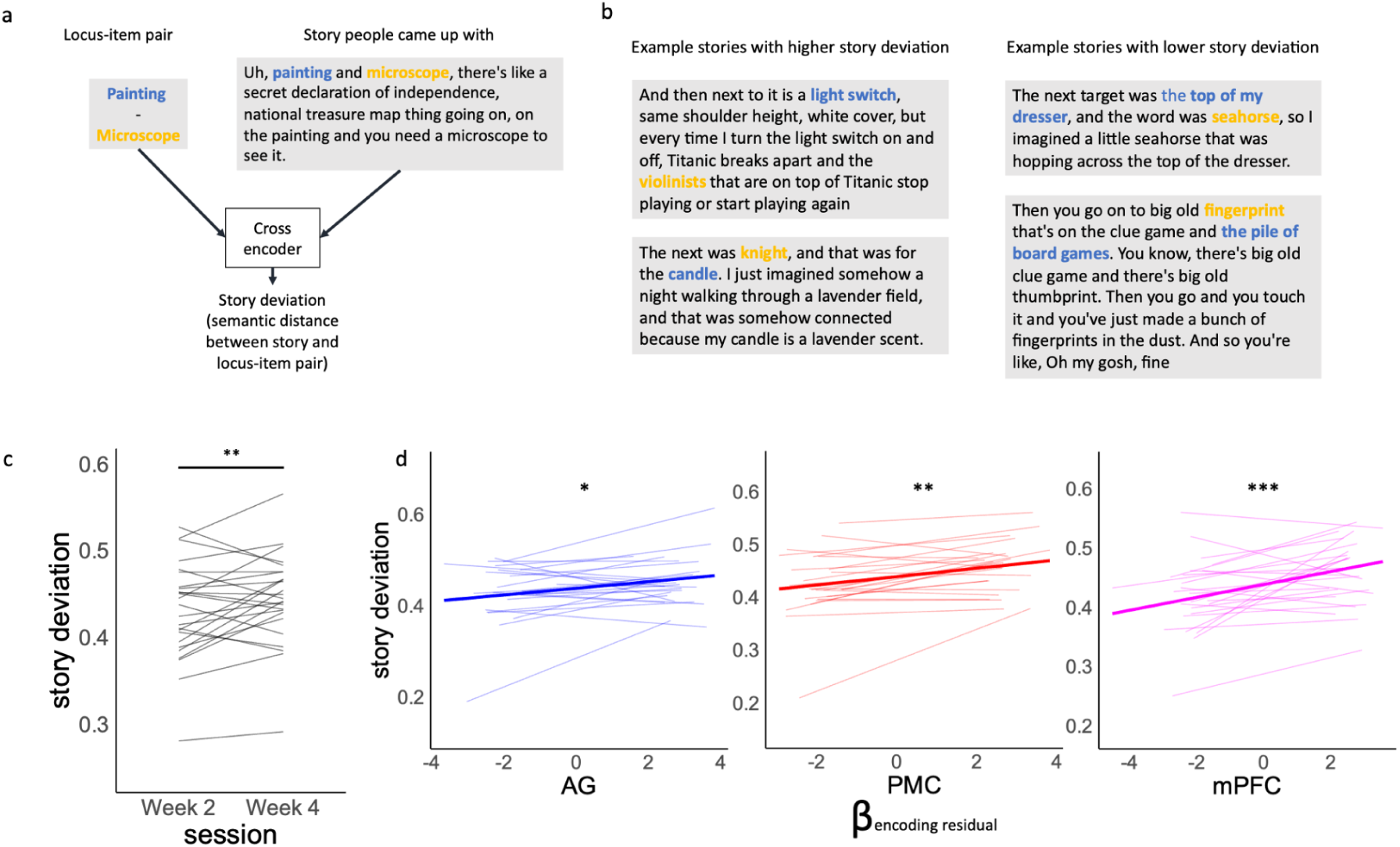
Measuring the novelty of generated stories and linking this measure to brain activity. **a.** How story deviation (semantic distance between the story and the locus-item pair) was measured using a cross-encoder language model. **b.** Example stories with high and low story deviation. In a and b, locus is highlighted in blue and item is highlighted in orange. **c.** How story deviation changes from week 2 to week 4. Each line represents a participant. **d.** Correlation between weight of encoding residual for a locus-item pair and story deviation for that pair. In all ROIs, the weight of encoding residual significantly predicted story deviation. Thick blue lines represent the overall trend and thin lines represent individual participants (in all three sessions). ( * p<0.05, ** p<0.01, *** p<0.001)

We next looked at whether brain activity measures (univariate activity, encoding residual weights) could predict the amount of the conjunctive representation in the verbal recalls of individual stories (operationalized using the story deviation measure described above). We first conducted mixed-effects linear regressions predicting the story deviation from univariate activity in each of the ROIs with random subject and session intercepts. In all three ROIs, univariate activity predicted higher story deviation values (AG: t = 4.29, p < .001; PMC: t = 3.40, p < .001; mPFC: t = 3.65, p < .001). Finally, we compared our *neural* conjunctivity measure (the weight of the encoding residual) to our *behavioral* conjunctivity measure (story deviation), conducting a linear mixed-effects regression predicting story deviation from the weight of encoding residual, with a random subject and session intercept (and controlling for duration of recall). We found that in all ROIs, a higher encoding residual weight for a locus-item pair predicted that the generated story for this pair would have a higher story deviation value (AG: t = 1.97, p = .049, PMC: t = 2.65, p = .008; mPFC: t = 3.35, p < .001). This analysis provides further evidence that novel patterns that are formed at encoding (not linearly related to locus or item patterns alone), and reinstated at retrieval, reflect the generation of new semantic details bridging between the item and locus.

## Discussion

In the current study, we trained a group of naive participants in the Method of Loci and used fMRI to measure the item-locus memories they formed during 3 sessions over the course of a month. Consistent with prior work (Wagner et al., 2021), all participants showed great improvement in their ability to remember word lists after learning MoL, demonstrating that the technique is highly effective and does not require users to have any exceptional baseline memory abilities. Our results highlight the role played by the mPFC and other DMN regions in representing schematic knowledge (loci) and items, both by themselves and in combination. Crucially, we found evidence that these regions contain conjunctive representations that track the content of the unique story details generated by each participant. The degree of neural conjunctive representation for locus-item pairs increased with training, and tracked the amount of novel details (going beyond the locus and item on their own) that were described by participants for each locus-item pair. Overall, these results point to a central role of conjunctive coding in the DMN for creating robust associative memories.

### Conjunctive representations in the DMN

Conjunctive representation involves combining stimulus elements into a representation that is more than the simple sum of its elements. In the past literature, forming conjunctive representations between arbitrary elements has been suggested to be one of the main functions of the hippocampus (McClelland et al., 1995; Rudy & Sutherland, 1995). In the current study, where people formed conjunctive representations between novel items and a well-established internal schema, we did not find evidence for the involvement of the hippocampus. Instead, we found widespread conjunctive representations in the cortex, particularly the DMN, when people connected episodic information (words) with a well-consolidated schema (their memory palace). We found that these regions, which have been previously implicated in processing schemas and forming memory consistent with schemas (e.g., Baldassano et al., 2018; Brod et al., 2015; Masís-Obando et al., 2021; Preston & Eichenbaum, 2013; Raykov et al., 2020, 2021; van Kesteren et al., 2012; Van Kesteren et al., 2013), play a key role in representing conjunctions between schemas and episodic details.

We found significant amounts of neural conjunctive representation in all three core regions of the DMN: AG, PMC, and mPFC. These regions have been previously implicated in work with memory for naturalistic events tied to familiar contexts (Chen et al., 2017; Zadbood et al., 2017), showing strong reinstatement of encoding patterns during retrieval. Our results suggest that conjunctive representations in these regions may also serve to “glue” schemas to event details when we form memories of naturalistic events (as in Baldassano et al., 2018; Masís-Obando et al., 2021). As noted in the introduction, it can be difficult to isolate the representations of schemas and event details in real world situations, which (in turn) makes it difficult to study how they are combined; the arbitrary pairings of details (words) and schemas (loci) in the current experiment allowed us to measure their separate and joint representations, providing unique insights into these processes in the brain.

In line with prior work, we found that mPFC played a central role in building schema-based memories, with significant effects across our analyses: the degree of conjunctive representation in mPFC robustly increased between week 2 and week 4; the neural similarity of conjunctive representations across pairs in mPFC tracked the semantic similarity of people’s stories; and the amount of mPFC conjunctive representation for individual pairs was strongly associated with the amount of conjunctive semantic detail in participants’ verbal recalls. Past lesion studies (Ghosh et al., 2014) have shown that vmPFC patients with confabulation symptoms had difficulty judging whether a word belonged to a script or not, suggesting that mPFC plays a role in relating current items to prior knowledge. Thus, mPFC may play an especially important role in the item-locus association step of MoL, drawing on prior knowledge to find a plausible interaction between the item and locus and elaborating on this interaction with new integrative details. For example, one participant (Figure 5b) linked a knight to a lavender candle by identifying the linking concept of a lavender field that the knight could walk through. This may draw on the same cognitive mechanisms as the “free generation of remote associates” task, in which participants attempt to generate a word with a meaningful but unusual relationship with a cue word, which is also known to be impaired by vmPFC lesions (Bendetowicz et al., 2018)

### Relating conjunctive coding to creativity and concept combination

In the current study, participants had complete freedom in how they chose to bind items to loci, and different participants came up with very different relationships even when provided with the same loci and items in the standardized-loci task. Although we were primarily interested in using MoL as a tool for studying memory, this experimental task could also be useful for creativity research. Creativity paradigms generally involve relatively constrained tasks like identifying a common word connecting two or three seemingly unrelated words (Bowden & Jung-Beeman, 2003) or finding alternative uses for a tool (Beaty et al., 2015), while our paradigm encouraged more open-ended divergent thinking (Runco & Yoruk, 2014) to generate semantic associates of the locus and/or the item. The current study thus provides a framework to study creativity across different semantic domains or across different individuals in a relatively unconstrained and spontaneous manner. The measurements developed in the study, including both the neural and semantic measure of conjunctive representation, could be relevant for providing insights into the neural mechanisms of creativity. Our finding that the amount of conjunctive representation in mPFC and PMC tracks the amount of semantic elaboration (as indexed by our story deviation measure) provides support for an important role of these regions in the creative process (Aziz-Zadeh et al., 2013; Shamay-Tsoory et al., 2011).

We hypothesize that MoL is relatively easy to learn (at least at a novice level) because conjunctively combining an item and locus draws on a fundamental cognitive function of building rich concepts from simpler primitives (e.g., Lake et al., 2015). While some of previous research has looked at the importance of the DMN in conceptual combination (Frankland & Greene, 2020), past research typically involved studying relatively straightforward combinations (like combining “old” and “woman”) and has focused more on the comprehension and perception of these ideas (Baron & Osherson, 2011). Our results argue for a role of the DMN in combining ideas together in a more elaborate way. While here we focus on how this conjunctive process supports episodic memory, future work can elucidate how conjunctive representations in the brain might support other processes such as concept learning (Zhou et al., 2024), episodic simulation and imagination (Spreng et al., 2018), and semantic-episodic linkage in trivia experts (Thieu et al., 2024).

One limitation of the current study is that, because we wanted to ensure that the novice participants would be able to form associations and that we would be able to accurately measure neural representations with fMRI, we provided 12 seconds for encoding each item. This is longer than most memory experiments with word list learning (e.g., Murdock, 1974) and other experiments involving MoL (e.g., Wagner et al., 2021), and is not how MoL is used by experts in a competition context, when an association needs to be formed in ∼2 sec. The long encoding duration was also one factor that led to performance being at ceiling even after just two weeks of training, making it infeasible to perform analyses relating conjunctivity to performance. Future work could examine whether very fast item-locus associations are supported by the same brain regions, as well as the meta-cognitive question of how mnemonists assess whether they have spent sufficient time adding conjunctive information for a specific association.

In conclusion, we used MoL as a window into studying the process of how people combine details with schemas in the context of memory formation. We found evidence of strong conjunctive representation in the DMN, which increased with training and tracked the semantic details added to the locus-item pair. These results demonstrate the importance of conjunctive representation in meaningfully combining schema with novel items and shed light on the neural mechanisms of creativity and concept combination.

## Methods

### Participants

26 novice participants passed our initial screening (see below), and were enrolled in a 4-week training program that included three fMRI scanning sessions. One participant was removed from the study for not demonstrating proper use of the MoL during training, resulting in a final sample of N=25. The participants’ age range from 18 to 48, with a mean age of 26.32 (SD = 7.46). The racial makeup of the participants were 16 White, 7 Asian, 1 Black or African American, and 1 mixed. Four of the participants were Hispanic or Latino. All participants were proficient in English. Participants were compensated $235 for the completion of the study. The experimental protocol was approved by the Institutional Review Board of Columbia University (AAAS0252).

### Stimuli

Participants self-generated 40 loci with the guidance of the coach, an expert user of MoL who led the training sessions. The loci were objects in a place familiar to the individual participants (e.g., light switch or kitchen stove in their room). In the first two scans, in the locus task, participants imagined and described these loci; in the encoding task, they used them to remember 40 words. The locus for a given trial was always generated internally by the participant based on their pre-practiced memory palace; they were not presented with the name / image of the loci during any of the fMRI tasks.

Participants also learned a standardized memory palace with 20 loci. The standardized memory palace consisted of five 2D virtual reality environments created in the game engine Unity. For each of the five environments, four distinct locations were selected, resulting in a total of 20 loci arranged in a fixed sequence. The order of the loci was the same for all participants. The standardized memory palace was taught to the participants through video clips created by rotating a virtual camera through the environments and stopping at each locus in the order.

Each locus was marked by a number (1 to 20) and an arrow pointing to it, along with a brief written description (e.g., “You turn the corner into the tavern and see an elevated table”, where elevated table is locus 1). 40 concrete nouns, 20 animate and 20 inanimate, were selected as the item stimuli used in all the scans and screening. These words were used in the item and encoding tasks in the first two scans, always appearing in a different random order. A subset of 20 words was selected and used in the item and encoding task in the last scan. In the item task the order of these 20 words was randomized, but in the encoding task, these words were presented to all the participants in the same order.

### Screening

Participants signed up through Columbia’s RecruitMe website, and went through a screening procedure, which consisted of a sequence of tasks similar to those used in the main fMRI experiment. They first completed two runs of the Item task, where a word was presented for 10 sec and participants reported whether the word was animate or inanimate during the final 5 sec of each trial. The purpose of including the item task was to better match the protocol used in the scanner, where two item localizers were conducted before completing the encoding task.

Participants then completed an encoding task, where each word was presented for 12 sec and they were instructed to remember the words in the right order. After that, participants attempted to retrieve the words one by one, in order. Participants then completed the fMRI safety form and a demographic form. Participants were selected based on availability for the 4-week memory training, and demographics (to ensure diversity in age and gender). After the first cohort of participants, we also selected participants based on memory performance in the screening part of the task (recalling fewer than 20 words in the right order). After participants were selected, they had a virtual meeting with the experimenter to complete the fMRI-related paperwork and schedule the fMRI scans.

### Training and scanning schedule

The experiment is conducted with groups of 3-5 people at a time. In the first two weeks, participants had four 1.5 hour interactive lectures with the coach, where they were trained to use the MoL by creating a personal memory palace with 40 anchors. After the training, they came in for the first fMRI scan (W2 scan). After the fMRI scan, participants went home and completed 10 daily practices. The daily practices consisted of an encoding and retrieval task that lasted 15 minutes each. They also meet with the coach one-on-one for check-in/questions before the next scan. They then came in for two consecutive days for two fMRI scans (W4D1, W4D2). After the W4D1 scan, they learned a standardized memory palace with 20 loci in it (as described above).

### Locus task

Participants were instructed to describe each locus in detail, and to keep talking until instructed to move to the next locus. Once the scan started, text appeared on the screen for 1 sec instructing the participants to start describing the first locus in their memory palace, then a fixation cross was shown on the center of the screen for 10 sec while participants gave their verbal description. After that, text appeared on the center of the screen for 1 sec instructing the participants to move to the next locus. This was repeated until all of the 40 or 20 loci were described.

### Item task

Participants were instructed to vividly imagine each word shown to them on the screen and make a judgment of whether the word is animate or inanimate. They were also instructed to not try to remember the words. Each word was shown to the participants for 10 sec, and 5 sec after the word was presented a text prompt appeared under the word asking participants to press button “1” if the word is animate and “2” if it is inanimate. After 10 sec, a fixation cross appeared on the center of the screen for 1 sec, followed by the next word.

### Encoding task

Participants were instructed to remember a list of words in order by using MoL. Each word was shown on the screen for 12 sec, followed by a fixation cross of 1 sec after each word.

### Retrieval task

After the scan started, participants were shown a fixation cross on the center of the screen. They were given unlimited time to recall all the words in order. They were told to talk about each locus and the item in detail, as well as how they associated the locus to the item. The recall was self-paced, but they were encouraged to spend at least 10 sec on each locus-item pair, even if they could not recall the item associated with a locus.

### Standardized memory palace learning and review

After the W4D1 scan, participants learned the 20 standardized loci by watching the videos of the standardized memory palace loci six times. In the first two repetitions (loci learning), participants were introduced to the loci one by one and had unlimited time to study each locus. They were told to press a button when they were ready to move on to the next locus. In the subsequent four repetitions (loci generation), participants viewed the video clips and text descriptions again but were prompted to type the name of the upcoming locus (e.g., elevated table) before advancing to the next one. After the six repetitions, participants were told to recall the names of all twenty loci in order. Before the W4D2 scan, participants reviewed the standardized memory palace by completing three rounds of loci generation. The experimenter then asked participants to verbally describe the 20 loci in order to ensure they could report all 20 without errors.

### Behavioral processing

For the locus and retrieval tasks, we used Open AI’s speech-to-text “Whisper” API (Radford et al., 2023) to obtain a transcript. Using the transcript and the audio recordings, we identified the locus described in each 10-sec time window in the locus task. For the retrieval task, we identified the locus-item pair (or in case participants forgot the word, we identified the locus spoken) and the start and end time when participants were describing each pair.

Participants wrote down their list of loci in order prior to the scan. Because participants were relatively new to the technique and their memory palace, they sometimes made mistakes in transitioning between their loci during the tasks. The most common was to skip a locus in either the locus, encoding, or retrieval task, and participants also occasionally misordered loci. To accommodate these errors, we used results from the retrieval task to (retrospectively) determine what locus was used at encoding. For example, consider the scenario where a participant’s loci were swing-grass-slide, and they were asked to remember the words apple-dog-pencil; in this case, if the participant recalled swing-apple, slide-dog, we assumed that they skipped a locus (grass) at encoding and they used slide to remember dog. Consequently, in the subsequent analyses we used the representation of the locus that was recalled (slide) to predict encoding/retrieval representation, rather than the locus that the participant should have used (grass).

### Performance scoring

To score participants’ performance in the memory task (which required participants to recall in the correct order), we identified the words recalled in the order they were spoken (repeated words were removed) and found the serial positions of these words from the encoding list. The word was considered to be recalled in the correct order if the serial position of the word was larger than the serial position of the previously recalled word.

### MRI Acquisition

Whole-brain data were acquired on a 3 Tesla Siemens Magnetom Prisma scanner equipped with a 64-channel head coil at Columbia University. Whole-brain, high-resolution (1.0 mm iso) T1 structural scans were acquired with a magnetization-prepared rapid acquisition gradient-echo sequence (MPRAGE) at the beginning of the scan session. Functional measurements were collected using a multiband echo-planar imaging (EPI) sequence (repetition time = 1.5s, echo time = 30ms, in-plane acceleration factor = 2, multiband acceleration factor = 3, voxel size = 2mm iso). Sixty-nine oblique axial slices were obtained in an interleaved order. All slices were tilted approximately −20 degrees relative to the AC-PC line. There were 6 functional runs in each scan: two runs of the locus task, two runs of item task, and one run of encoding task, and one run of retrieval task.

### fMRI preprocessing

Results included in this manuscript come from preprocessing performed using fMRIPrep 23.0.2 (Esteban et al. (2019); Esteban et al. (2018); RRID:SCR_016216), which is based on Nipype 1.8.6 (Esteban et al., (2022); Gorgolewski et al., (2011); RRID:SCR_002502).

### Anatomical data preprocessing

A total of 1 T1-weighted (T1w) images were found within the input BIDS dataset.The T1-weighted (T1w) image was corrected for intensity non-uniformity (INU) with N4BiasFieldCorrection (Tustison et al., 2010), distributed with ANTs 2.3.3 (Avants et al., 2008, RRID:SCR_004757), and used as T1w-reference throughout the workflow. The T1w-reference was then skull-stripped with a Nipype implementation of the antsBrainExtraction.sh workflow (from ANTs), using OASIS30ANTs as target template. Brain tissue segmentation of cerebrospinal fluid (CSF), white-matter (WM) and gray-matter (GM) was performed on the brain-extracted T1w using fast (FSL 6.0.5.1:57b01774, RRID:SCR_002823,(Zhang, Brady, and Smith, 2001). Brain surfaces were reconstructed using recon-all (FreeSurfer 7.3.2, RRID:SCR_001847, (Dale, Fischi, Sereno, 1999), and the brain mask estimated previously was refined with a custom variation of the method to reconcile ANTs-derived and FreeSurfer-derived segmentations of the cortical gray-matter of Mindboggle (RRID:SCR_002438, Klein et al. 2017). Volume-based spatial normalization to one standard space (MNI152NLin2009cAsym) was performed through nonlinear registration with antsRegistration (ANTs 2.3.3), using brain-extracted versions of both T1w reference and the T1w template. The following template was were selected for spatial normalization and accessed with TemplateFlow (23.0.0, Ciric et al., 2022): ICBM 152 Nonlinear Asymmetrical template version 2009c [Fonov et al., (2009), RRID:SCR_008796; TemplateFlow ID: MNI152NLin2009cAsym].

### Preprocessing of B0 inhomogeneity mappings

A total of 3 fieldmaps were found available within the input BIDS structure for this particular subject. A deformation field to correct for susceptibility distortions was estimated based on fMRIPrep’s fieldmap-less approach. The deformation field is that resulting from co-registering the EPI reference to the same-subject T1w-reference with its intensity inverted (Wang et al., 2017; Huntenburg, 2014). Registration is performed with antsRegistration (ANTs 2.3.3), and the process regularized by constraining deformation to be nonzero only along the phase-encoding direction, and modulated with an average fieldmap template (Treiber et al., 2016).

### Functional data preprocessing

For each of the 18 BOLD runs found per subject (across all tasks and sessions), the following preprocessing was performed. First, a reference volume and its skull-stripped version were generated using a custom methodology of fMRIPrep. Head-motion parameters with respect to the BOLD reference (transformation matrices, and six corresponding rotation and translation parameters) are estimated before any spatiotemporal filtering using mcflirt (FSL 6.0.5.1:57b01774, Jenkinson et al., 2002). The estimated fieldmap was then aligned with rigid-registration to the target EPI (echo-planar imaging) reference run. The field coefficients were mapped on to the reference EPI using the transform. The BOLD reference was then co-registered to the T1w reference using bbregister (FreeSurfer) which implements boundary-based registration (Greve & Fischl, 2009). Co-registration was configured with six degrees of freedom. Several confounding time-series were calculated based on the preprocessed BOLD: framewise displacement (FD), DVARS and three region-wise global signals. FD was computed using two formulations following Power (absolute sum of relative motions, Power et al., (2014)) and Jenkinson (relative root mean square displacement between affines, Jenkinson et al., (2002)). FD and DVARS are calculated for each functional run, both using their implementations in Nipype (following the definitions by Power et al., 2014). The three global signals are extracted within the CSF, the WM, and the whole-brain masks. Additionally, a set of physiological regressors were extracted to allow for component-based noise correction (CompCor, Behzadi et al., 2007). Principal components are estimated after high-pass filtering the preprocessed BOLD time-series (using a discrete cosine filter with 128s cut-off) for the two CompCor variants: temporal (tCompCor) and anatomical (aCompCor). tCompCor components are then calculated from the top 2% variable voxels within the brain mask. For aCompCor, three probabilistic masks (CSF, WM and combined CSF+WM) are generated in anatomical space.

The implementation differs from that of Behzadi et al. in that instead of eroding the masks by 2 pixels on BOLD space, a mask of pixels that likely contain a volume fraction of GM is subtracted from the aCompCor masks. This mask is obtained by dilating a GM mask extracted from the FreeSurfer’s aseg segmentation, and it ensures components are not extracted from voxels containing a minimal fraction of GM. Finally, these masks are resampled into BOLD space and binarized by thresholding at 0.99 (as in the original implementation). Components are also calculated separately within the WM and CSF masks. For each CompCor decomposition, the k components with the largest singular values are retained, such that the retained components’ time series are sufficient to explain 50 percent of variance across the nuisance mask (CSF, WM, combined, or temporal). The remaining components are dropped from consideration. The head-motion estimates calculated in the correction step were also placed within the corresponding confounds file. The confound time series derived from head motion estimates and global signals were expanded with the inclusion of temporal derivatives and quadratic terms for each (Satterthwaite et al., 2013). Frames that exceeded a threshold of 0.5 mm FD or 1.5 standardized DVARS were annotated as motion outliers. Additional nuisance timeseries are calculated by means of principal components analysis of the signal found within a thin band (crown) of voxels around the edge of the brain, as proposed by (Patriat Reynolds, and Birn, 2017). The BOLD time-series were resampled into standard space, generating a preprocessed BOLD run in MNI152NLin2009cAsym space. First, a reference volume and its skull-stripped version were generated using a custom methodology of fMRIPrep. The BOLD time-series were resampled onto the following surfaces (FreeSurfer reconstruction nomenclature): fsaverage6.

All resamplings can be performed with a single interpolation step by composing all the pertinent transformations (i.e. head-motion transform matrices, susceptibility distortion correction when available, and co-registrations to anatomical and output spaces). Gridded (volumetric) resamplings were performed using antsApplyTransforms (ANTs), configured with Lanczos interpolation to minimize the smoothing effects of other kernels (Lanczos, 1964). Non-gridded (surface) resamplings were performed using mri_vol2surf (FreeSurfer).

Many internal operations of fMRIPrep use Nilearn 0.9.1 (Abraham et al., 2014, RRID:SCR_001362), mostly within the functional processing workflow. For more details of the pipeline, see the section corresponding to workflows in fMRIPrep’s documentation.

### ROI and searchlight definition

We used ROIs in the default mode network previously found to be responsive to schematic content (Baldassano et al., 2018): angular gyrus (1868 vertices), medial prefrontal cortex (mPFC; 2069 vertices), and posterior medial cortex (PMC; 2495 vertices). These ROIs were originally derived from a resting-state network atlas on the fsaverage6 surface (Thomas Yeo et al., 2011).

Searchlight ROIs were defined as circular regions on the cortical surface, by identifying all vertices within 11 edges of a center vertex along the fsaverage6 mesh. Since the average edge length between vertices is 1.4mm, searchlights had a radius of approximately 15mm. We defined a circular searchlight around every vertex on a hemisphere, and then iteratively removed the most redundant searchlights (i.e. those whose vertices were covered by the most other searchlights). We stopped removing searchlights when doing so would cause some vertices to be covered by fewer than six searchlights. This yielded approximately 1000 searchlights on each hemisphere.

### Activity pattern extraction

After fMRIPrep, the data (now in fsaverage6 and MNI152 space) were further preprocessed by a custom python script that removed from the data (via linear regression) any variance related to the six degrees of freedom motion correction estimate and their derivatives, mean signals in the CSF and white matter, motion outlier timepoints (defined above), and a cosine basis set for high-pass filtering w/ 0.008 Hz (125s) cut-off. We then z scored each run to have zero mean and standard deviation of 1. All subsequent analyses, described below, were performed using custom python and R scripts.

To obtain the locus, item and encoding patterns, we conducted a GLM predicting whole-brain univariate activity from the design matrices for the corresponding tasks based on the timing of each locus/item. For the retrieval task, the design matrix was based on the manual identification of the start and end time of describing each locus-item pair. The coefficients from fitting this GLM were used as the values for defining the voxel patterns of the locus/item/pair.

### Pattern similarity analyses: Locus and item representations (Figure 2)

To look at where loci were represented in the brain, for the loci that were described in both locus runs, we correlated the representations of the pairs of loci between the two runs for each participant in each session. This produced a correlation matrix of the representational similarity between each locus in run 1 with each locus in run 2 for each searchlight. We then computed the representational similarity of the same loci across two runs (the average of the diagonal of the correlation matrix). For assessing statistical significance, the similarity between different loci (the mean off-diagonal of the correlation matrix) was then subtracted from the mean similarity of the same loci. Once we obtained the difference, we randomly shuffled the rows of the correlation matrix 1000 times to compute a null distribution of this diagonal vs off-diagonal difference, and computed the z-score for the searchlight

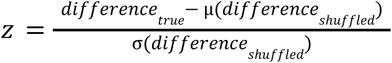

which represents the degree to which there was a locus-specific representation across runs. A z value was computed for each surface vertex as the average of z values from all searchlights that included that vertex. We then combined maps across all subjects and all sessions, and ran a one-sample t-test of the z values against 0. The r values in the voxels with q < .05 after false discovery rate (FDR) correction from the one-sample t-test were plotted in the map.

For the locus-encoding and item-encoding similarity analysis, the approach was the same as described above. Here, we looked at the similarity between the average of representations of the two locus/item runs and the encoding representation where the locus/item is used. If participants only described a locus in one of the locus runs, then the representation of the locus from the single run was used.

### Relating the similarity of encoding residuals to the similarity of stories (Figure 3)

First, for each locus-item pair on week 4 day 2 (separately for each participant and each brain region), we obtained the residual of the encoding representation by regressing out the representations of the locus and item that were used during encoding. Then, separately for each brain region, we generated a *neural correlation matrix* for each item-locus pair by computing the Pearson’s correlation of each participant’s encoding residual with each other participant’s encoding residual for that pair. Next, we took the verbal stories that were generated on week 4 day 2 for each locus-item pair, and we generated a *semantic correlation matrix* for that pair using the cross-encoder version of the Sentence-BERT language model (Reimers & Gurevych, 2019). This model takes two text passages as input and generates a score (0 - 1) quantifying the semantic similarity between the two passages. Then the lower triangles of the neural and semantic correlation matrices for all the locus-item pairs were each flattened into a long vector. Finally, a Spearman correlation was computed between the neural and semantic correlation vector for each pair. A permutation test was conducted, shuffling the semantic vectors between locus-item pairs 1000 times to obtain a chance level neural-semantic correlation. We averaged across the 20 locus-item pairs in each permutation, generating 1000 null correlations. For each ROI, the significance level was determined by counting the percentage of times that the 1000 random correlations were more extreme than the true correlation.

### Identifying conjunctive representations by predicting retrieval representations (Figure 4)

For each item-locus pair (separately for each participant and searchlight), we ran a GLM predicting the representation of that locus-item pair at retrieval from the locus by itself, the item by itself, and the encoding residual representation for the locus-item pair; this analysis incorporated data from all of the sessions. To make sure the weights reflected item-specific reinstatement rather than general task-related patterns, we adjusted the weights against a null distribution in which retrieval of each locus-item pair was predicted by the locus, item, and encoding residual representations for a different locus-item pair. The labels for the retrieval representations within a session were shuffled 100 times. For each locus-item pair, we calculated the adjusted weight, β_*task*_:

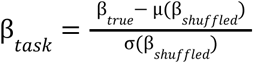

The weights were averaged across all the pairs from each session for each participant for each searchlight. A β value was computed for each surface vertex as the average of β values from all searchlights that included that vertex. For each vertex, a one-sample t-test compared the participant-level average weights (across all three sessions) against 0. To compare the difference in amount of representation between encoding residual and locus/item, we conducted a paired t-test on the difference in average weights of encoding residual and locus/item within participants in each session in each participant.

We also ran the above analysis for the three ROIs; here, we compared β_*task*_ against 0 for each ROI for each timepoint (week 2 or 4) using a one-sample t-test. For across-region comparisons, we conducted paired t-tests between each pair of regions, combining week 2 and week 4 data. For looking at differences between week 2 and 4 in the amount of conjunctive representation, we conducted a linear mixed effect regression. We controlled for speaking duration as a nuisance regressor to ensure that differences between weeks were not being driven solely by longer recall durations for each item (which provide more timepoints for estimating retrieval patterns and therefore less-noisy representations). We used the following formula:

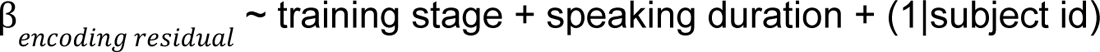

### Semantic conjunctive representation (Figure 5)

We conceptualized semantic conjunctive representation as the amount of additional details added to the locus and item pairs. To measure this, we computed *story deviation*, the semantic distance between the locus-item pair on its own and the whole story, using the same cross-encoder model described above. After we obtained the similarity score, we subtract the score from 1 to generate the story deviation measure. To look at changes in story deviation, we conducted a linear mixed effect regression with the following formula:

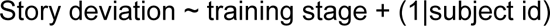

For each ROI, we looked at how univariate activity during encoding is related to story deviation, with a linear mixed effects regression:

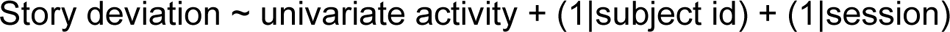

We also looked at how weights of encoding residuals are related to story deviation, with a linear mixed effects regression:

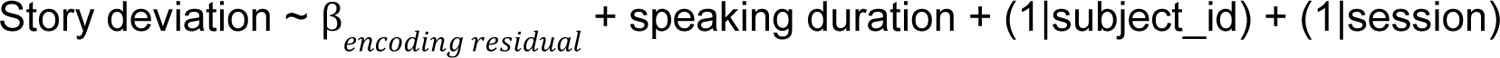

---

**Figure S1.**
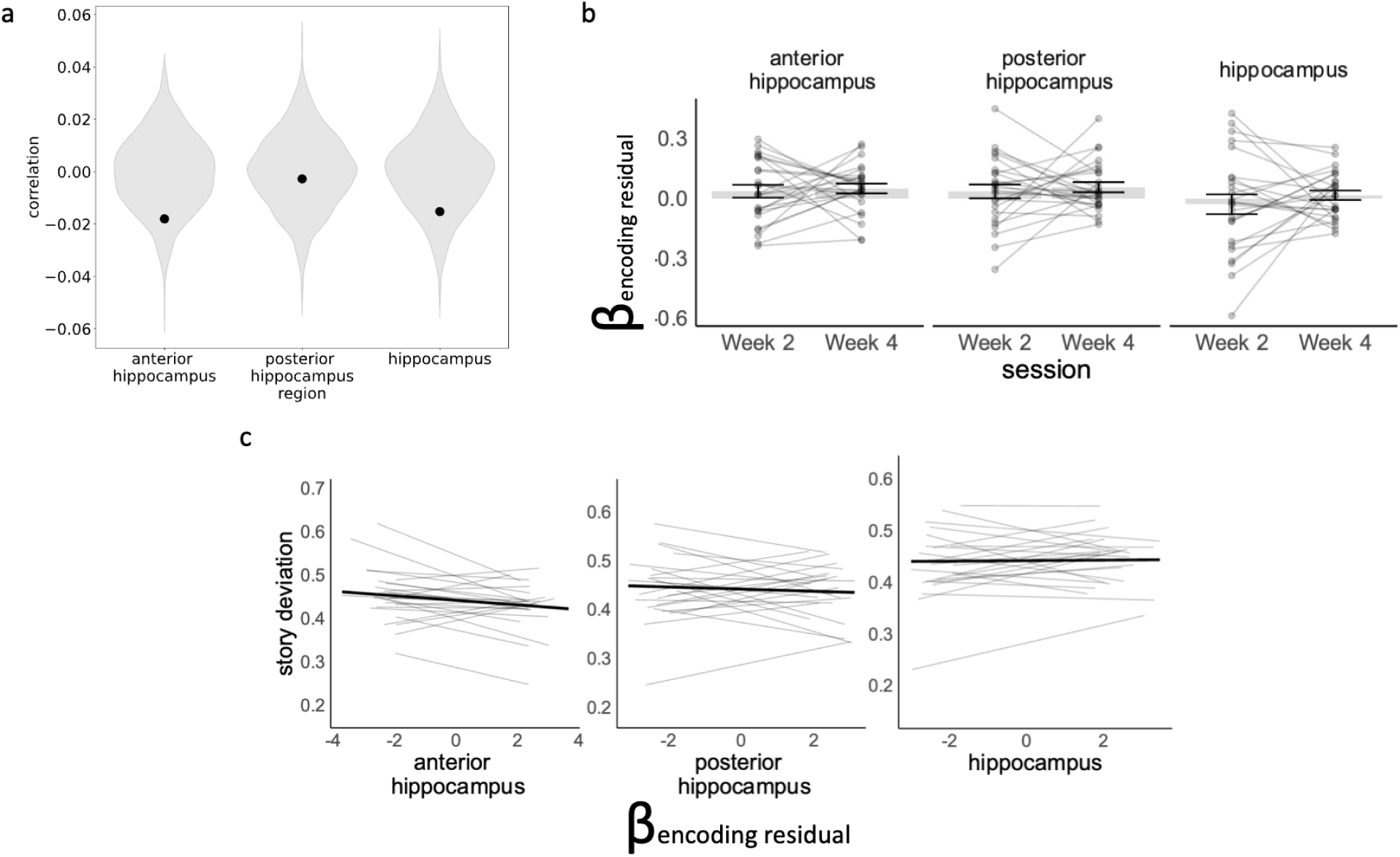
Results of the ROI analyses done in hippocampal ROIs (anterior, posterior, and whole). a. There was no evidence that encoding residual in any of the hippocampal ROIs tracked semantic similarity in the story told by the participants. b. There was no evidence for changes in weights of encoding residual in hippocampus over the course of the training, despite the anterior and posterior hippocampus showing weights significantly above zero at week 4 (anterior: t = 2.07, p = .05, posterior: t = 2.25, p = .034). c. There was no evidence that the amount of encoding residual is related to story deviation.

## References

10.1016/j.cognition.2023.105711

Abraham, A., Pedregosa, F., Eickenberg, M., Gervais, P., Mueller, A., Kossaifi, J., Gramfort, A., Thirion, B., & Varoquaux, G. (2014). Machine learning for neuroimaging with scikit-learn. Frontiers in Neuroinformatics, 8. 10.3389/fninf.2014.00014

Anderson, J. R. (1981). Effects of prior knowledge on memory for new information. Memory & Cognition, 9(3), 237–246. 10.3758/BF03196958

Anderson, R. C., & Pichert, J. W. (1978). Recall of previously unrecallable information following a shift in perspective. Journal of Verbal Learning and Verbal Behavior, 17(1), 1–12. 10.1016/S0022-5371(78)90485-1

Avants, B., Epstein, C., Grossman, M., & Gee, J. (2008). Symmetric diffeomorphic image registration with cross-correlation: Evaluating automated labeling of elderly and neurodegenerative brain. Medical Image Analysis, 12(1), 26–41. 10.1016/j.media.2007.06.004

Aziz-Zadeh, L., Liew, S.-L., & Dandekar, F. (2013). Exploring the neural correlates of visual creativity. Social Cognitive and Affective Neuroscience, 8(4), 475–480. 10.1093/scan/nss021

Baldassano, C., Beck, D. M., & Fei-Fei, L. (2017). Human–Object Interactions Are More than the Sum of Their Parts. Cerebral Cortex, 27(3), 2276–2288. 10.1093/cercor/bhw077

Baldassano, C., Hasson, U., & Norman, K. A. (2018). Representation of Real-World Event Schemas during Narrative Perception. Journal of Neuroscience, 38(45), 9689–9699. 10.1523/JNEUROSCI.0251-18.2018

Baron, S. G., & Osherson, D. (2011). Evidence for conceptual combination in the left anterior temporal lobe. NeuroImage, 55(4), 1847–1852. 10.1016/j.neuroimage.2011.01.066

Bartlett, S. F. C. (1932). Remembering: A Study in Experimental and Social Psychology. Cambridge University Press.

Beaty, R. E., Benedek, M., Barry Kaufman, S., & Silvia, P. J. (2015). Default and Executive Network Coupling Supports Creative Idea Production. Scientific Reports, 5(1), 10964. 10.1038/srep10964

Behzadi, Y., Restom, K., Liau, J., & Liu, T. T. (2007). A component based noise correction method (CompCor) for BOLD and perfusion based fMRI. NeuroImage, 37(1), 90–101. 10.1016/j.neuroimage.2007.04.042

Bellezza, F. S. (1996). Chapter 10—Mnemonic Methods to Enhance Storage and Retrieval. In E. L. Bjork & R. A. Bjork (Eds.), Memory (pp. 345–380). Academic Press. 10.1016/B978-012102570-0/50012-4

Bendetowicz, D., Urbanski, M., Garcin, B., Foulon, C., Levy, R., Bréchemier, M.-L., Rosso, C., Thiebaut de Schotten, M., & Volle, E. (2018). Two critical brain networks for generation and combination of remote associations. Brain, 141(1), 217–233. 10.1093/brain/awx294

Bowden, E. M., & Jung-Beeman, M. (2003). Aha! Insight experience correlates with solution activation in the right hemisphere. Psychonomic Bulletin & Review, 10(3), 730–737. 10.3758/BF03196539

Brod, G., Lindenberger, U., Werkle-Bergner, M., & Shing, Y. L. (2015). Differences in the neural signature of remembering schema-congruent and schema-incongruent events. NeuroImage, 117, 358–366. 10.1016/j.neuroimage.2015.05.086

Chase, W. G., & Simon, H. A. (1973). Perception in chess. Cognitive Psychology, 4(1), 55–81. 10.1016/0010-0285(73)90004-2

Chen, J., Leong, Y. C., Honey, C. J., Yong, C. H., Norman, K. A., & Hasson, U. (2017). Shared memories reveal shared structure in neural activity across individuals. Nature Neuroscience, 20(1), 115–125. 10.1038/nn.4450

Ciric, R., Thompson, W. H., Lorenz, R., Goncalves, M., MacNicol, E. E., Markiewicz, C. J., Halchenko, Y. O., Ghosh, S. S., Gorgolewski, K. J., Poldrack, R. A., & Esteban, O. (2022). TemplateFlow: FAIR-sharing of multi-scale, multi-species brain models. Nature Methods, 19(12), 1568–1571. 10.1038/s41592-022-01681-2

Dale, A. M., Fischl, B., & Sereno, M. I. (1999). Cortical Surface-Based Analysis. NeuroImage, 9(2), 179–194. 10.1006/nimg.1998.0395

De Soares, A., Kim, T., Mugisho, F., Zhu, E., Lin, A., Zheng, C., & Baldassano, C. (2024). Top-down attention shifts behavioral and neural event boundaries in narratives with overlapping event scripts. Current Biology, 34(20), 4729–4742.e5. 10.1016/j.cub.2024.09.013

Eich, E. (1985). Context, memory, and integrated item/context imagery. *Journal of Experimental Psychology: Learning*, Memory, and Cognition, 11(4), 764–770. 10.1037/0278-7393.11.1-4.764

Erez, J., Cusack, R., Kendall, W., & Barense, M. D. (2016). Conjunctive Coding of Complex Object Features. Cerebral Cortex, 26(5), 2271–2282. 10.1093/cercor/bhv081

Esteban, O., Markiewicz, C. J., Blair, R. W., Moodie, C. A., Isik, A. I., Erramuzpe, A., Kent, J. D., Goncalves, M., DuPre, E., Snyder, M., Oya, H., Ghosh, S. S., Wright, J., Durnez, J., Poldrack, R. A., & Gorgolewski, K. J. (2019). fMRIPrep: A robust preprocessing pipeline for functional MRI. Nature Methods, 16(1), 111–116. 10.1038/s41592-018-0235-4

Esteban, O., Markiewicz, C. J., Burns, C., Goncalves, M., Jarecka, D., Ziegler, E., Berleant, S., Ellis, D. G., Pinsard, B., Madison, C., Waskom, M., Notter, M. P., Clark, D., Manhães-Savio, A., Clark, D., Jordan, K., Dayan, M., Halchenko, Y. O., Loney, F., … Ghosh, S. (2022). nipy/nipype: 1.8.3 (Version 1.8.3) [Computer software]. Zenodo. 10.5281/ZENODO.596855

Esteban, O., Markiewicz, C. J., Goncalves, M., Provins, C., Kent, J. D., DuPre, E., Salo, T., Ciric, R., Pinsard, B., Blair, R. W., Poldrack, R. A., & Gorgolewski, K. J. (2018). fMRIPrep: A robust preprocessing pipeline for functional MRI (Version 23.0.2) [Computer software]. Zenodo. 10.5281/zenodo.7863421

Fonov, V., Evans, A., McKinstry, R., Almli, C., & Collins, D. (2009). Unbiased nonlinear average age-appropriate brain templates from birth to adulthood. NeuroImage, 47, S102. 10.1016/S1053-8119(09)70884-5

Frankland, S. M., & Greene, J. D. (2020). Concepts and Compositionality: In Search of the Brain’s Language of Thought. Annual Review of Psychology, 71(Volume 71, 2020), 273–303. 10.1146/annurev-psych-122216-011829

Gasser, C., & Davachi, L. (2023). Cross-Modal Facilitation of Episodic Memory by Sequential Action Execution. Psychological Science, 34(5), 581–602. 10.1177/09567976231158292

Ghosh, V. E., & Gilboa, A. (2014). What is a memory schema? A historical perspective on current neuroscience literature. Neuropsychologia, 53, 104–114. 10.1016/j.neuropsychologia.2013.11.010

Ghosh, V. E., Moscovitch, M., Melo Colella, B., & Gilboa, A. (2014). Schema Representation in Patients with Ventromedial PFC Lesions. The Journal of Neuroscience, 34(36), 12057–12070. 10.1523/JNEUROSCI.0740-14.2014

Gilboa, A., & Marlatte, H. (2017). Neurobiology of Schemas and Schema-Mediated Memory. Trends in Cognitive Sciences, 21(8), 618–631. 10.1016/j.tics.2017.04.013

Gobet, F., & Waters, A. J. (2003). The Role of Constraints in Expert Memory. *Journal of Experimental Psychology: Learning*, Memory, and Cognition, 29(6), 1082–1094. 10.1037/0278-7393.29.6.1082

Gorgolewski, K., Burns, C. D., Madison, C., Clark, D., Halchenko, Y. O., Waskom, M. L., & Ghosh, S. S. (2011). Nipype: A Flexible, Lightweight and Extensible Neuroimaging Data Processing Framework in Python. Frontiers in Neuroinformatics, 5. 10.3389/fninf.2011.00013

Greve, D. N., & Fischl, B. (2009). Accurate and robust brain image alignment using boundary-based registration. NeuroImage, 48(1), 63–72. 10.1016/j.neuroimage.2009.06.060

Howard, M. W., & Kahana, M. J. (2002). A Distributed Representation of Temporal Context. Journal of Mathematical Psychology, 46(3), 269–299. 10.1006/jmps.2001.1388

Huang, J., Furness, E., Liu, Y., Kenmoe, M.-J., Elias, R., Zeng, H. T., & Baldassano, C. (2023). Accurate predictions facilitate robust memory encoding independently from stimulus probability. OSF. 10.31234/osf.io/dhzvc

Huang, J., Velarde, I., Ma, W. J., & Baldassano, C. (2023). Schema-based predictive eye movements support sequential memory encoding. eLife, 12, e82599. 10.7554/eLife.82599

Huntenburg, J. (2014). Evaluating nonlinear coregistration of BOLD EPI and T1w images. Master’s Thesis, Berlin: Freie Universität. http://hdl.handle.net/11858/00-001M-0000-002B-1CB5-A.

Jenkinson, M., Bannister, P., Brady, M., & Smith, S. (2002). Improved Optimization for the Robust and Accurate Linear Registration and Motion Correction of Brain Images. NeuroImage, 17(2), 825–841. 10.1006/nimg.2002.1132

Kriegeskorte, N., Mur, M., & Bandettini, P. (2008). Representational similarity analysis—Connecting the branches of systems neuroscience. Frontiers in Systems Neuroscience, 2. https://www.frontiersin.org/article/10.3389/neuro.06.004.2008

Lake, B. M., Salakhutdinov, R., & Tenenbaum, J. B. (2015). Human-level concept learning through probabilistic program induction. Science, 350(6266), 1332–1338. 10.1126/science.aab3050

Lanczos, C. (1964). Evaluation of Noisy Data. Journal of the Society for Industrial and Applied Mathematics Series B Numerical Analysis, 1(1), 76–85. 10.1137/0701007

Liang, J. C., Erez, J., Zhang, F., Cusack, R., & Barense, M. D. (2020). Experience Transforms Conjunctive Object Representations: Neural Evidence for Unitization After Visual Expertise. Cerebral Cortex, 30(5), 2721–2739. 10.1093/cercor/bhz250

Liu, C., Ye, Z., Chen, C., Axmacher, N., & Xue, G. (2022). Hippocampal Representations of Event Structure and Temporal Context during Episodic Temporal Order Memory. Cerebral Cortex, 32(7), 1520–1534. 10.1093/cercor/bhab304

Liu, Z.-X., Grady, C., & Moscovitch, M. (2017). Effects of Prior-Knowledge on Brain Activation and Connectivity During Associative Memory Encoding. Cerebral Cortex, 27(3), 1991–2009. 10.1093/cercor/bhw047

Maguire, E. A., Valentine, E. R., Wilding, J. M., & Kapur, N. (2003). Routes to remembering: The brains behind superior memory. Nature Neuroscience, 6(1), 90–95. 10.1038/nn988

Masís-Obando, R., Norman, K. A., & Baldassano, C. (2021). Schema representations in distinct brain networks support narrative memory during encoding and retrieval (p. 2021.05.17.444363). 10.1101/2021.05.17.444363

Masís-Obando, R., Norman, K. A., & Baldassano, C. (2024). How sturdy is your memory palace? Reliable room representations predict subsequent reinstatement of placed objects (p. 2024.11.26.625465). bioRxiv. 10.1101/2024.11.26.625465

McClelland, J. L., McNaughton, B. L., & O’Reilly, R. C. (1995). Why there are complementary learning systems in the hippocampus and neocortex: Insights from the successes and failures of connectionist models of learning and memory. Psychological Review, 102(3), 419–457. 10.1037/0033-295X.102.3.419

Morris, R. G. M. (2006). Elements of a neurobiological theory of hippocampal function: The role of synaptic plasticity, synaptic tagging and schemas. European Journal of Neuroscience, 23(11), 2829–2846. 10.1111/j.1460-9568.2006.04888.x

Murdock, B. B. (1974). Human memory: Theory and data (pp. x, 362). Lawrence Erlbaum.

Murnane, K., Phelps, M. P., & Malmberg, K. (1999). Context-dependent recognition memory: The ICE theory. Journal of Experimental Psychology: General, 128(4), 403–415. 10.1037/0096-3445.128.4.403

O’Reilly, R. C., & Rudy, J. W. (2001). Conjunctive representations in learning and memory: Principles of cortical and hippocampal function. Psychological Review, 108(2), 311–345. 10.1037/0033-295X.108.2.311

Patriat, R., Reynolds, R. C., & Birn, R. M. (2017). An improved model of motion-related signal changes in fMRI. NeuroImage, 144, 74–82. 10.1016/j.neuroimage.2016.08.051

Polyn, S. M., Norman, K. A., & Kahana, M. J. (2009). A context maintenance and retrieval model of organizational processes in free recall. Psychological Review, 116(1), 129–156. 10.1037/a0014420

Power, J. D., Mitra, A., Laumann, T. O., Snyder, A. Z., Schlaggar, B. L., & Petersen, S. E. (2014). Methods to detect, characterize, and remove motion artifact in resting state fMRI. NeuroImage, 84, 320–341. 10.1016/j.neuroimage.2013.08.048

Preston, A. R., & Eichenbaum, H. (2013). Interplay of Hippocampus and Prefrontal Cortex in Memory. Current Biology, 23(17), R764–R773. 10.1016/j.cub.2013.05.041

Puff, C. R. (1979). Memory organization and structure. Academic Press. https://cir.nii.ac.jp/crid/1130000797374336256

Radford, A., Kim, J. W., Xu, T., Brockman, G., Mcleavey, C., & Sutskever, I. (2023). Robust Speech Recognition via Large-Scale Weak Supervision. Proceedings of the 40th International Conference on Machine Learning, 28492–28518. https://proceedings.mlr.press/v202/radford23a.html

Raykov, P. P., Keidel, J. L., Oakhill, J., & Bird, C. M. (2020). The brain regions supporting schema-related processing of people’s identities. Cognitive Neuropsychology, 37(1–2), 8–24. 10.1080/02643294.2019.1685958

Raykov, P. P., Keidel, J. L., Oakhill, J., & Bird, C. M. (2021). Activation of Person Knowledge in Medial Prefrontal Cortex during the Encoding of New Lifelike Events. *Cerebral Cortex*, bhab027. 10.1093/cercor/bhab027

Reagh, Z. M., & Ranganath, C. (2023). Flexible reuse of cortico-hippocampal representations during encoding and recall of naturalistic events. Nature Communications, 14(1), Article 1. 10.1038/s41467-023-36805-5

Reimers, N., & Gurevych, I. (2019). *Sentence-BERT: Sentence Embeddings using Siamese BERT-Networks* (arXiv:1908.10084). arXiv. 10.48550/arXiv.1908.10084

Rudy, J. W., & Sutherland, R. J. (1995). Configural association theory and the hippocampal formation: An appraisal and reconfiguration. Hippocampus, 5(5), 375–389. 10.1002/hipo.450050502

Runco, M. A., & Yoruk, S. (2014). The Neuroscience of Divergent Thinking. Activitas Nervosa Superior, 56(1), 1–16. 10.1007/BF03379602

Satterthwaite, T. D., Elliott, M. A., Gerraty, R. T., Ruparel, K., Loughead, J., Calkins, M. E., Eickhoff, S. B., Hakonarson, H., Gur, R. C., Gur, R. E., & Wolf, D. H. (2013). An improved framework for confound regression and filtering for control of motion artifact in the preprocessing of resting-state functional connectivity data. NeuroImage, 64, 240–256. 10.1016/j.neuroimage.2012.08.052

Shamay-Tsoory, S. G., Adler, N., Aharon-Peretz, J., Perry, D., & Mayseless, N. (2011). The origins of originality: The neural bases of creative thinking and originality. Neuropsychologia, 49(2), 178–185. 10.1016/j.neuropsychologia.2010.11.020

Shin, Y. S., Masís-Obando, R., Keshavarzian, N., Dáve, R., & Norman, K. A. (2021). Context-dependent memory effects in two immersive virtual reality environments: On Mars and underwater. Psychonomic Bulletin & Review, 28(2), 574–582. 10.3758/s13423-020-01835-3

Spreng, R. N., Madore, K. P., & Schacter, D. L. (2018). Better imagined: Neural correlates of the episodic simulation boost to prospective memory performance. Neuropsychologia, 113, 22–28. 10.1016/j.neuropsychologia.2018.03.025

Squire, L. R., Shimamura, A. P., & Amaral, D. G. (1989). 12—Memory and the Hippocampus. In J. H. Byrne & W. O. Berry (Eds.), Neural Models of Plasticity (pp. 208–239). Academic Press. 10.1016/B978-0-12-148955-7.50016-3

Thieu, M. K., Wilkins, L. J., & Aly, M. (2024). Episodic-semantic linkage for $1000: New semantic knowledge is more strongly coupled with episodic memory in trivia experts. Psychonomic Bulletin & Review, 31(4), 1867–1879. 10.3758/s13423-024-02469-5

Thomas Yeo, B. T., Krienen, F. M., Sepulcre, J., Sabuncu, M. R., Lashkari, D., Hollinshead, M., Roffman, J. L., Smoller, J. W., Zöllei, L., Polimeni, J. R., Fischl, B., Liu, H., & Buckner, R. L. (2011). The organization of the human cerebral cortex estimated by intrinsic functional connectivity. Journal of Neurophysiology, 106(3), 1125–1165. 10.1152/jn.00338.2011

Treiber, J. M., White, N. S., Steed, T. C., Bartsch, H., Holland, D., Farid, N., McDonald, C. R., Carter, B. S., Dale, A. M., & Chen, C. C. (2016). Characterization and Correction of Geometric Distortions in 814 Diffusion Weighted Images. PLOS ONE, 11(3), e0152472. 10.1371/journal.pone.0152472

Tse, D., Langston, R. F., Kakeyama, M., Bethus, I., Spooner, P. A., Wood, E. R., Witter, M. P., & Morris, R. G. M. (2007). Schemas and Memory Consolidation. Science, 316(5821), 76–82. 10.1126/science.1135935

Tustison, N. J., Avants, B. B., Cook, P. A., Yuanjie Zheng, Egan, A., Yushkevich, P. A., & Gee, J. C. (2010). N4ITK: Improved N3 Bias Correction. IEEE Transactions on Medical Imaging, 29(6), 1310–1320. 10.1109/TMI.2010.2046908

van den Honert, R. N., McCarthy, G., & Johnson, M. K. (2017). Holistic versus feature-based binding in the medial temporal lobe. Cortex, 91, 56–66. 10.1016/j.cortex.2017.01.011

Van Kesteren, M. T. R., Beul, S. F., Takashima, A., Henson, R. N., Ruiter, D. J., & Fernández, G. (2013). Differential roles for medial prefrontal and medial temporal cortices in schema-dependent encoding: From congruent to incongruent. Neuropsychologia, 51(12), 2352–2359. 10.1016/j.neuropsychologia.2013.05.027

van Kesteren, M. T. R., Rignanese, P., Gianferrara, P. G., Krabbendam, L., & Meeter, M. (2020). Congruency and reactivation aid memory integration through reinstatement of prior knowledge. Scientific Reports, 10(1), 4776. 10.1038/s41598-020-61737-1

van Kesteren, M. T. R., Ruiter, D. J., Fernández, G., & Henson, R. N. (2012). How schema and novelty augment memory formation. Trends in Neurosciences, 35(4), 211–219. 10.1016/j.tins.2012.02.001

Wagner, I. C., Konrad, B. N., Schuster, P., Weisig, S., Repantis, D., Ohla, K., Kühn, S., Fernández, G., Steiger, A., Lamm, C., Czisch, M., & Dresler, M. (2021). Durable memories and efficient neural coding through mnemonic training using the method of loci. Science Advances, 7(10), eabc7606. 10.1126/sciadv.abc7606

Wang, S., Peterson, D. J., Gatenby, J. C., Li, W., Grabowski, T. J., & Madhyastha, T. M. (2017). Evaluation of Field Map and Nonlinear Registration Methods for Correction of Susceptibility Artifacts in Diffusion MRI. Frontiers in Neuroinformatics, 11. 10.3389/fninf.2017.00017

Watkins, M. J., & Gardiner, J. M. (1979). An appreciation of generate-recognize theory of recall. Journal of Verbal Learning and Verbal Behavior, 18(6), 687–704. 10.1016/S0022-5371(79)90397-9

Zadbood, A., Chen, J., Leong, Y. C., Norman, K. A., & Hasson, U. (2017). How We Transmit Memories to Other Brains: Constructing Shared Neural Representations Via Communication. *Cerebral Cortex (New York*, N.Y.: 1991), *27*(10), 4988–5000. 10.1093/cercor/bhx202

Zhang, Y., Brady, M., & Smith, S. (2001). Segmentation of brain MR images through a hidden Markov random field model and the expectation-maximization algorithm. IEEE Transactions on Medical Imaging, 20(1), 45–57. 10.1109/42.906424

Zhou, Y., Feinman, R., & Lake, B. M. (2024). Compositional diversity in visual concept learning. Cognition, 244, 105711. 10.1016/j.cognition.2023.105711

